# Mechanisms and fluid dynamics of foraging in heterotrophic nanoflagellates

**DOI:** 10.1101/2021.04.01.438049

**Authors:** Sei Suzuki, Anders Andersen, Thomas Kiørboe

## Abstract

Heterotrophic nanoflagellates are the main consumers of bacteria and picophytoplankton in the ocean. In their micro-scale world, viscosity impedes predator-prey contact, and the mechanisms that allow flagellates to daily clear a volume of water for prey corresponding to 10^6^ times their own volume is unclear. It is also unclear what limits observed maximum ingestion rates of about 10^4^ bacterial prey per day. We used high-speed video-microscopy to describe feeding flows, flagellum kinematics, and prey searching, capture, and handling in four species with different foraging strategies. In three species, prey-handling times limit ingestion rates and account well for their reported maximum values. Similarly, observed feeding flows match reported clearance rates. Simple point-force models allowed us to estimate the forces required to generate the feeding flows, between 4-13 pN, and consistent with the force produced by the hairy (hispid) flagellum, as estimated using resistive force theory. Hispid flagella can produce a force that is much higher than the force produced by a naked flagellum with similar kinematics, and the hairy flagellum is therefore key to foraging in most nanoflagellates. Our findings provide a mechanistic underpinning of observed functional responses of prey ingestion rates in nanoflagellates.

## INTRODUCTION

Heterotrophic nanoflagellates play a key role in microbial food webs in the oceans by feeding on phytoplankton and bacteria and by transferring primary production to higher trophic levels when. Their top-down control shapes the structure and function of microbial communities and mediate essential biogeochemical cycles in the sea [1–4]. Despite their importance, the mechanisms of prey capture and the processes limiting their ingestion rates are not fully understood [5, 6].

Flagellates live in a low Reynolds number world where viscosity impedes predator-prey contact [7]. Yet, nanoflagellates are capable of daily clearing a volume of water for prey that corresponds to about one million times their cell volume [8, 9]. In the nutritionally dilute ocean, this is the clearance rate needed to sustain a viable population in the face of predation mortality [10]. How the flagellates overcome the impeding effect of viscosity is unclear for many forms.

Most flagellates use their flagella to swim, to generate feeding currents, and to capture prey. Many studies have examined the fluid dynamics of flagellates from the perspective of swimming, but few have done so from the perspective of food acquisition [11–16], even though feeding is likely a more fundamental component of the fitness than propulsion *per se*. In a few cases, the flagellum forces have been estimated indirectly from swimming speeds [13] or from quantification of feeding flows [16, 17]. However, there is large variation in flagellar kinematics and arrangements between species that yields big differences in the strength and architecture of the feeding flows [18]. In most cases, the forces generated by the flagellum and required to account for the necessary high clearance rates are unknown.

Direct observations of flagellate feeding were pioneered by Sleigh and Fenchel [43, 24], and followed by few additional studies [16, 20–25]. These studies revealed a variety of prey acquisition and handling strategies. Prey is either intercepted by the cell body, a flagellum, or specialized structures, and then either rejected or transported to the spot on the cell surface where it is phagocytized. During capture and handling of prey, the feeding current may cease, and no further prey can be captured [20, 25]. Handling time may therefore put an upper limit on prey ingestion rate. The maximum clearance rate governed by the feeding current, and the maximum ingestion rate, potentially governed by the prey handling time, together describe the functional response of the prey ingestion rate as function of prey concentration. This is the key function characterizing predator-prey interactions.

The aim of this study is to provide a mechanistic underpinning of the functional response relations that have been obtained in incubation experiments [8, 9]. We build on and expand previous observational work on flagellate foraging, and we describe predator-prey encounters and prey handling in four species with characteristic predation modes. We portray the flagellar dynamics during the different grazing phases and quantify prey handling times to evaluate the potential for prey ingestion. We further quantify the feeding flow, estimate clearance rates from observed flow fields, and use simple fluid dynamic models to compute the forces needed to account for the observed flows as well as the forces that the flagellum produces.

## RESULTS

### Prey capture and handling

Supplemetary movies 1-4 illustrate the different behaviors described below; and morphometric data and flagellum properties can be found in Supplementary Tab. S1.

*Paraphysomonas foraminifera* (Fig. 1) attaches to the surface by a filamentous structure from the posterior end of the cell. Cells are located directly on the surface or at a distance. On the anterior side there are two flagella, both with their base near the ingestion site. When searching for prey, the long flagellum continuously beats in a curved fashion (46 ± 6 Hz) and creates a feeding current towards the cell, while the second shorter flagellum is inactive. When a food particle enters the feeding current, it is pulled towards the flagellate (Fig. 2a). The flagellate responds to the prey before it establishes visible contact with the flagellum (Fig. 2b). Most likely the first contact is with the invisible flagellar hairs. As also observed by Christensen-Dalsgaard and Fenchel, the presence of prey is followed by a series of changes in flagellar behavior [13]. The end of the long flagellum hooks over into a fixed position while the wave amplitude and the beating frequency increases (67 ± 8 Hz) and the short flagellum starts beating rapidly (104 ± 15 Hz). The particle is transported longitudinally until it is confined between the two flagella (Fig. 2c). Finally, the prey is positioned between the short flagellum and the body, ready for phagocytosis (Fig. 2d). During ingestion, three possible scenarios were observed. In the first case, the long flagellum returns to its original position and beating frequency; thus a feeding current is generated immediately (Fig. 2e). Alternatively, the long flagellum returns to the searching position but with a reduced beating frequency (28 ± 7 Hz after 2 seconds); therefore the flow rate is not restored until after more than 2 s. A third scenario involves an immobilized, stiff and wavy long flagellum whilst the short flagellum continued beating until finally pausing. The flagellate remained inactive for a long period, which usually exceeded the recording capacity. Off-the-record observations confirmed that after these long breaks, *P*. *foraminifera* starts beating again to search for more prey. To reject a captured particle, the flagellate releases the prey by returning the long flagellum to the original beating pattern and position (Fig. 2f), and continues creating a feeding current (Fig. 2g).

**Figure 1.**
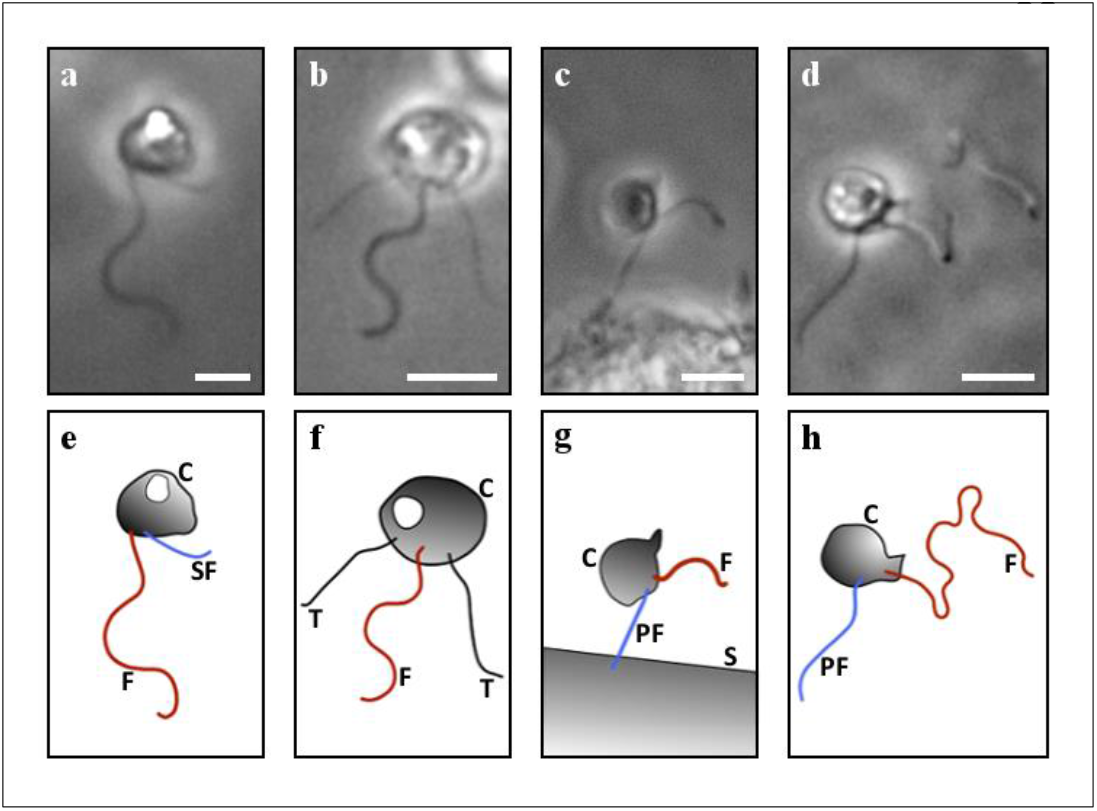
Phase contrast micrographs of the studied flagellates *Paraphysomonas foraminifera* (a and e), *Pteridomonas danica* (b and f), *Cafeteria roenbergensis* (c and g), and *Pseudobodo* sp. (d and h). Abbreviations in drawings (e, f, g, h): C – cell, F – flagellum, SF – short flagellum, T – tentacles, PF – posterior flagellum, and S – surface. Scale bar = 5 μm.

**Figure 2.**
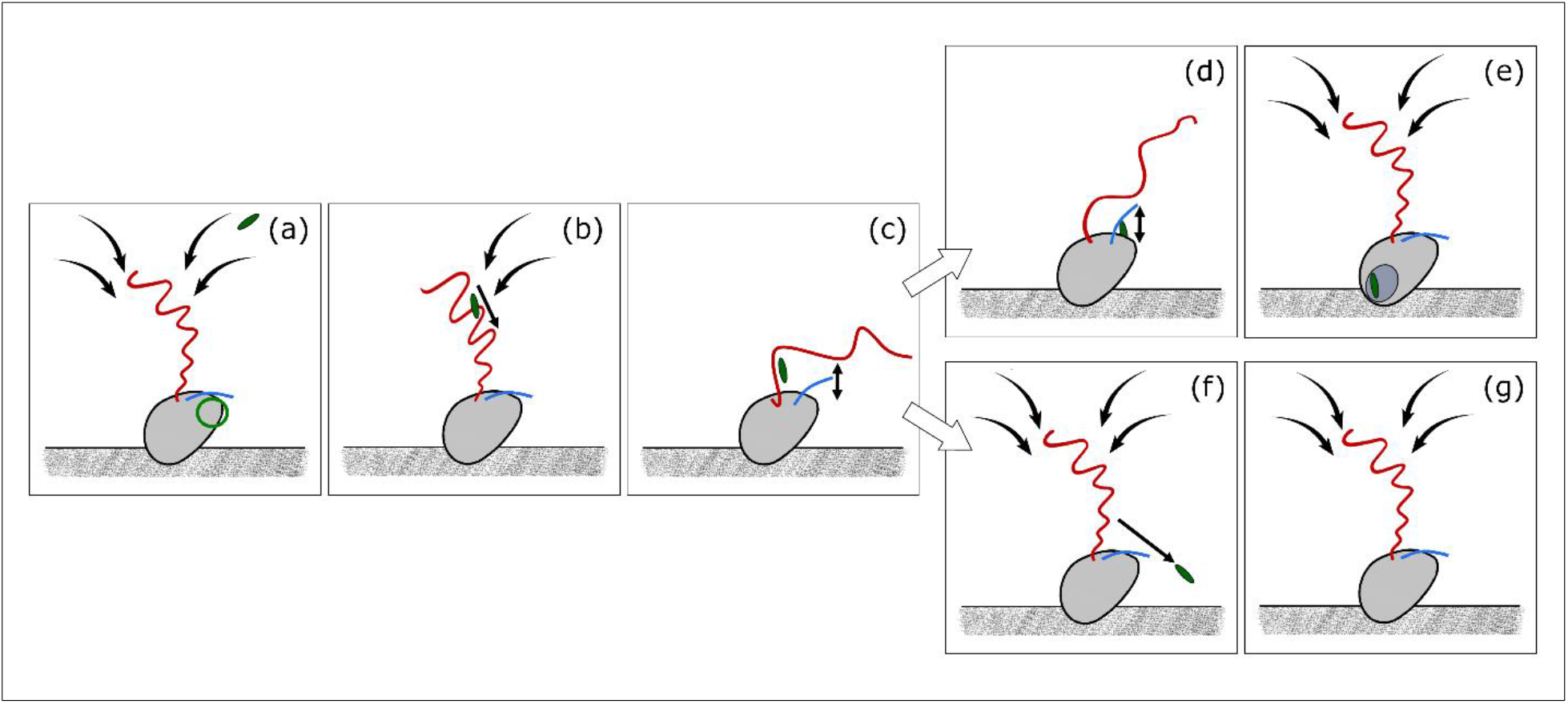
Schematic representation of foraging by *Paraphysomonas foraminifera*. Prey handling steps: searching and capture (a - c); prey ingestion (d -e); and rejection (f - g). Figure objects: long and curved flagellum (red), short flagellum (blue), ingestion site (green circle), feeding current (solid curved arrows), object in motion (solid straight arrows).

*Ochromonas moestrupii* and *Chrysophyceae* have a similar feeding behavior as *P*. *foraminifera*. The prevailing difference is their straight, long, beating flagellum (52 ± 9 Hz and 50 ± 5 Hz, respectively) in contrast to the curved flagellum that characterizes *P*. *foraminifera*. All three species attach posteriorly in the same manner; and contact and handle the prey with comparable flagellar behaviors for ingestions and rejections.

The handling time of *P*. *foraminifera* starts when the prey establishes contact with the flagellum. Rejected prey are handled more quickly than ingested prey (Fig. 3 and Supplementary Tab. S2). Handling times were independent of prey size over the range of encountered prey sizes (Supplementary Fig. S1). When the flagellate paralyzed during an ingestion, the handling time ended when the flagellum reactivated, and the feeding current was restored. This frozen flagellum behavior was reported in 6 out of the 20 ingestions, which 5 exceeded the remaining recording time of 9 – 45 s. Thus these 5 observations were excluded from further analysis due to their unknown duration.

**Figure 3.**
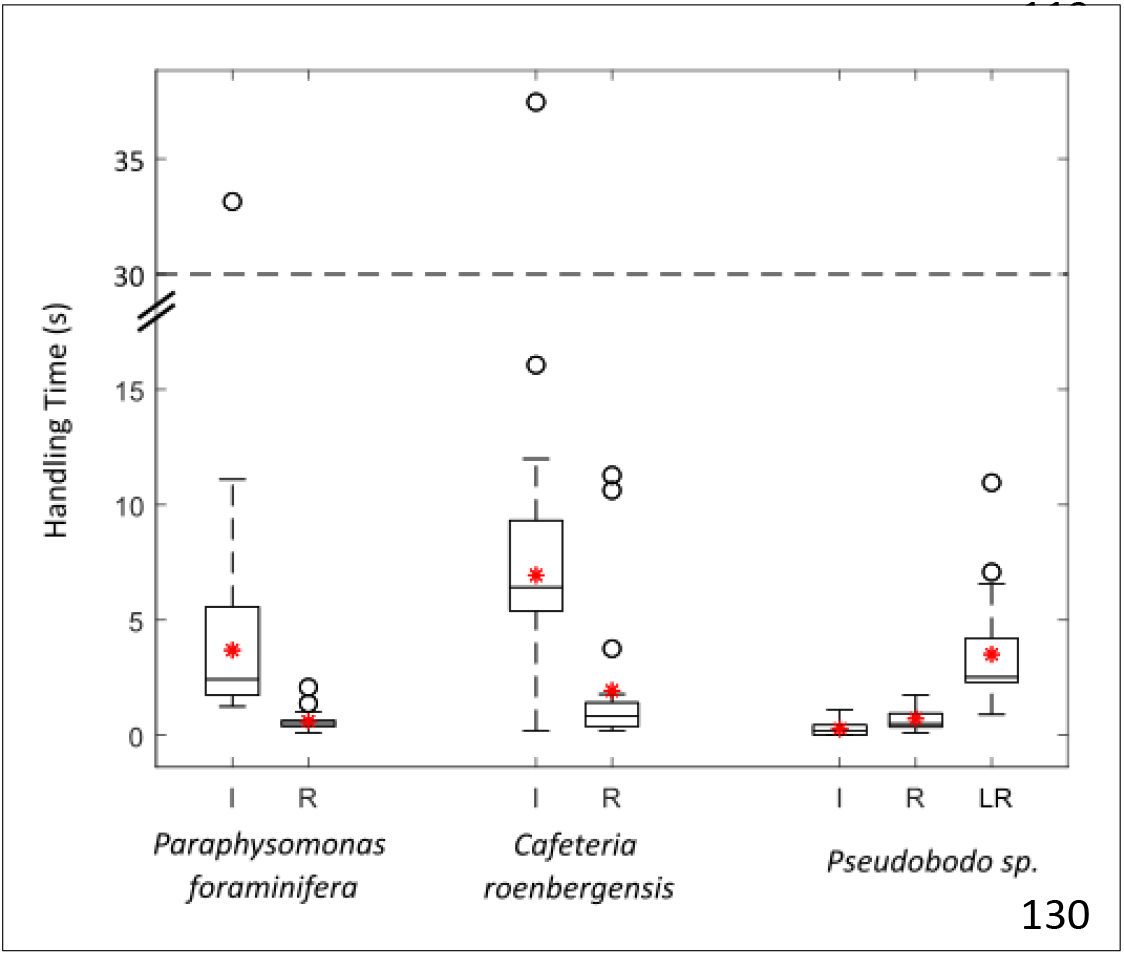
Handling times by *P. foraminifera*\*, *C. roenbergensis*, and *Pseudobodo* sp. Boxplots for the durations of ingestions (I), rejections (R), and lash rejections (LR); from prey capture to resuming the feeding current (median – dividing black line in the boxplot; mean – asterisk; error bars – dashed lines; outliers – empty circles). * Not including the ingestion cases of *P. foraminifera* that were limited by the recording time (N = 5).

*Pteridomonas danica* (Fig. 1) is attached to the surface with a posterior stalk. Its flagellum beats in a plane with constant frequency, creating a current towards the cell and perpendicular to the attachment surface as also described by Christensen-Dalsgaard and Fenchel [13]. Prey arriving in the flow is intercepted by the tentacle ‘crown’. When food is captured by the tentacles, it is slowly transported towards the cell for phagocytosis. Some particles move outwards and accumulate at the tentacle tips before drifting away. It is unclear if this is an active rejection or a result of the local flow. The beating pattern of the flagellate remains uniform throughout all prey encounters, behaving purely like a filtering predator. More than one food particle can be captured or handled simultaneously, and the handling time for *P. danica* is therefore zero.

When sessile, *Cafeteria roenbergensis* (Fig. 1) attaches to the surface with the tip of the posterior flagellum that flexes at irregular intervals. The anterior flagellum beats with constant frequency in a three-dimensional pattern with separate power and recovery phases to create a slightly erratic feeding current parallel to the attachment surface (Fig. 4a). As previously observed [25], prey particles are intercepted by the cell, not the flagellum (Fig. 4b). Upon prey contact on the sensitive frontal side of the predator, the anterior flagellum stops beating and rapidly arches fully extended against the prey. Thus, food is physically retained between the flagellum and the cell, close to where phagocytosis takes place (Fig. 4c). If the prey establishes first contact elsewhere on the cell, the flagellum continues beating while the food is transported along the cell surface, upstream towards the frontal area. When the prey gets near the ingestion site, the flagellum stops beating to capture the prey and initiate phagocytosis (Fig. 4d). The anterior flagellum resumes its initial beating behavior while or after the prey is phagocytized (Fig. 4e). The flagellate can reject captured prey by returning the flagellum to its original position and releasing the food (Fig. 4f and 4g). Handling times for *C. roenbergensis* are shorter for rejected than for ingested prey (Fig. 3), and durations were uncorrelated with the prey size (Supplementary Fig. S1).

**Figure 4.**
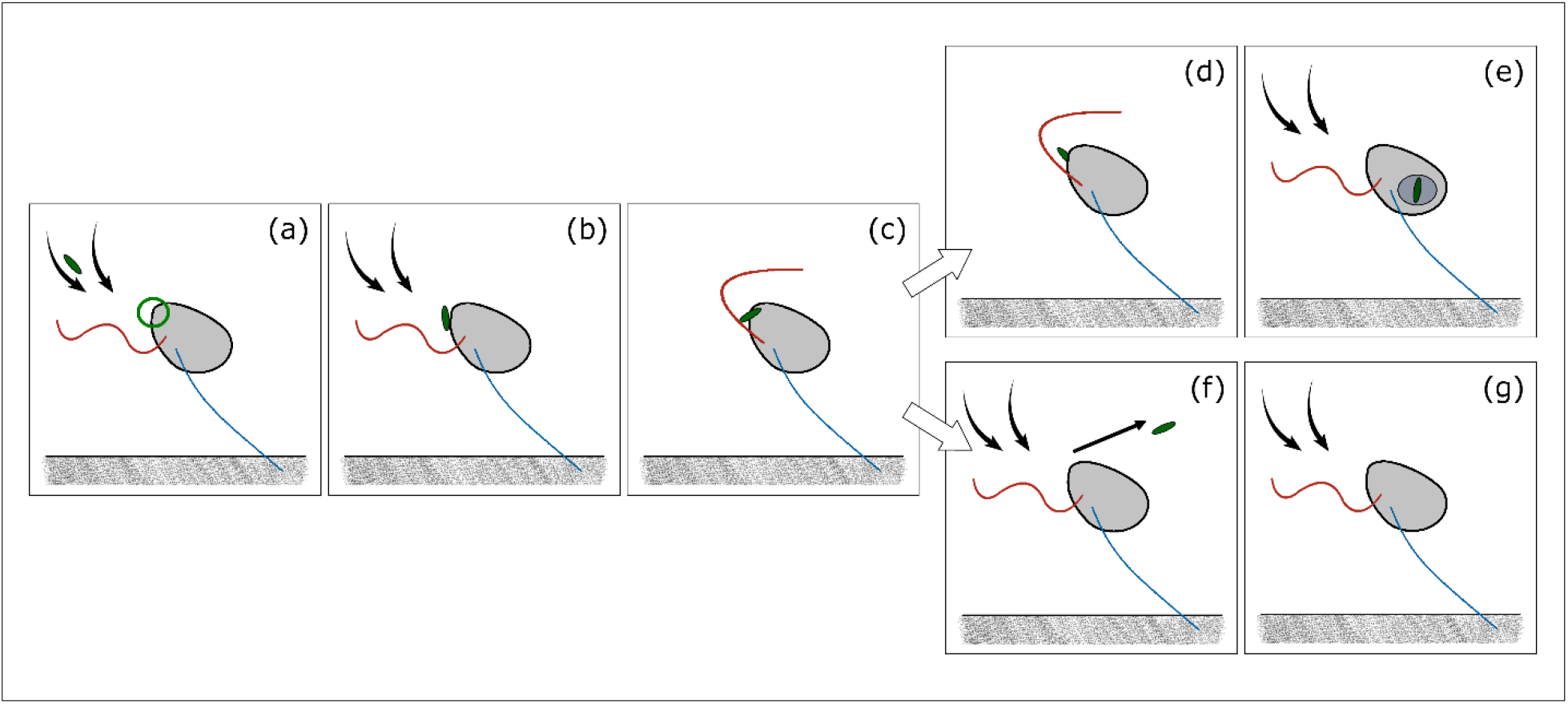
Schematic representation of foraging by *Cafeteria roenbergensis*. Prey handling steps: searching and capture (a - c); prey ingestion (d -e); and rejection (f - g). Figure objects: frontal flagellum (red), posterior flagellum (blue), ingestion site (green circle), feeding current (solid curved arrows), object in motion (solid straight arrows).

When free-swimming, *Pseudobodo* sp. (Fig. 1) is pulled forwards by the extended, beating anterior flagellum while the posterior flagellum inactively trails behind. *Pseudobodo* sp. attach to surfaces while feeding with the long anterior flagellum resembling a lasso loop (Fig. 5a). The flagellum beats (36 ± 10 Hz) and creates a feeding flow through the loop as briefly described previously [19]. The flow direction can vary from parallel to perpendicular to the surface; and distance between loop and cell (5.7 ± 2.8 μm) is variable during the searching mode. Food particles are intercepted by the anterior flagellum (Fig. 5b). Prey contact triggers an increase in beating frequency (63 ± 15 Hz) at a reduced wave amplitude, and a shorter helical pitch. The prey is retained between the cell and the flagellum and transported towards the body (Fig. 5c), either for ingestion (Fig. 5d-e) or rejection. *Pseudobodo* sp. has two ways to actively reject a particle: 1) the quick release and 2) the lash rejection. While the particle is captured between the flagellum and the body, it can be quickly released by reducing the beating frequency and returning the flagellum to its original position (Fig. 5f). Then, the feeding flow is rapidly restored (Fig. 5g). In the lash rejection, the flagellum stops beating for an instant before starting to ‘uncoil’ from base to end, sometimes finalizing fully extended and straight (Fig. 5h). Then it slowly starts beating (6 ± 3 Hz) with a higher wavelength and amplitude (2.2 ± 0.4 μm). In this rejection mode the prey is physically pushed away by the flagellate after being captured. Once the prey is released, the flagellum slowly coils back and recovers the initial ‘loop’ beating pattern (Fig. 5i). *Pseudobodo* sp. rejected particles with diameter smaller than 3 μm with a quick release or a lash rejection, in contrast to particles with diameter larger than 5 μm that were only discriminated with the later strategy (Supplementary Fig. S1). The handling times of captured particles in *Pseudobodo* sp. were rather short and of similar duration for particles ingested or released quickly, while the lash rejections were of much longer duration (Fig. 3). For almost half of the ingested particles (8/20) the flagellate intercepted and processed the prey without modifying the flagellar beat, thus the handling time was zero. Similar to the other species, handling times were independent of prey size (Supplementary Fig. S1).

**Figure 5.**
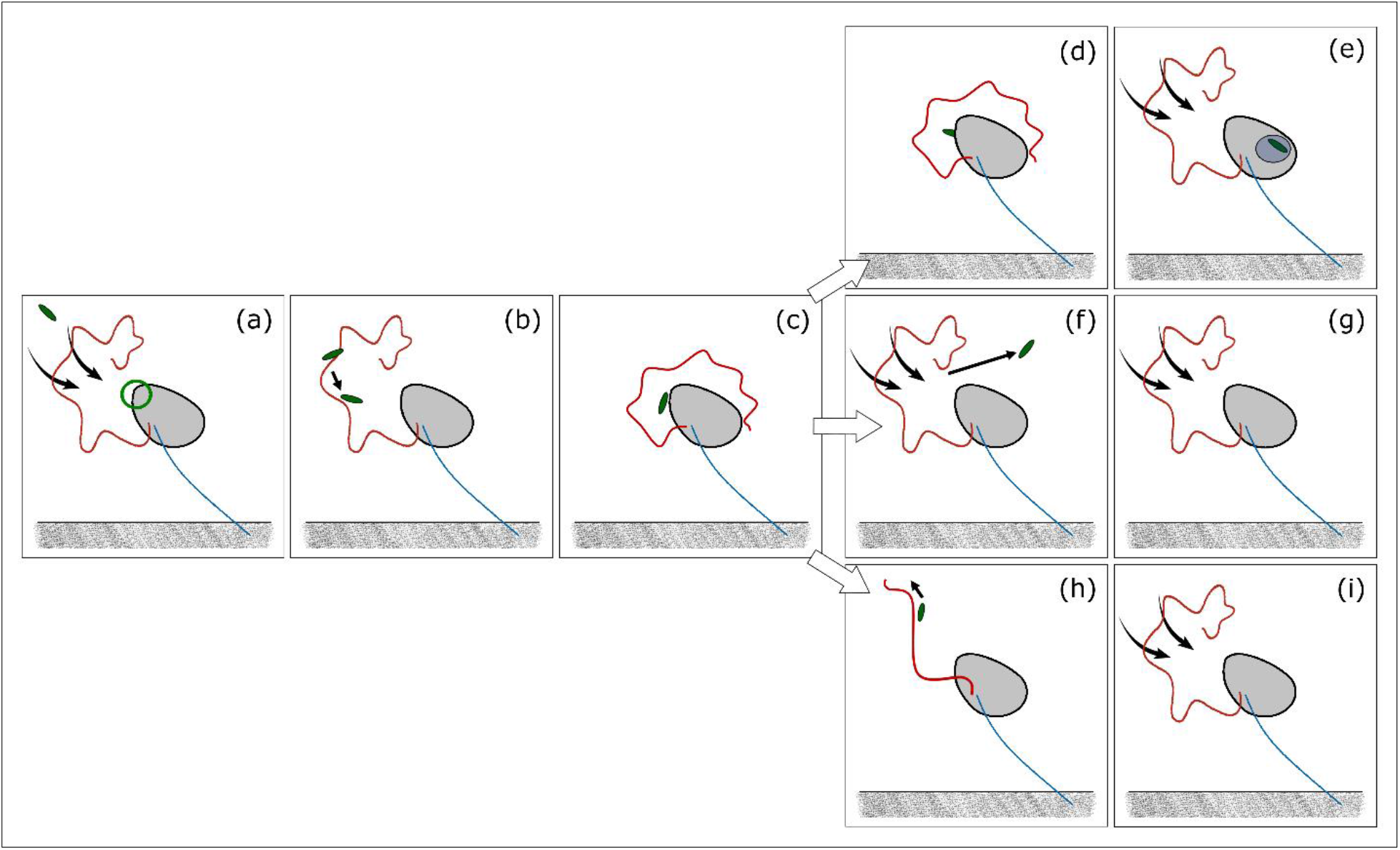
Schematic representation of foraging by *Pseudobodo* sp. Prey handling steps: searching and capture (a - c); prey ingestion (d -e); rejection (f - g); and ‘lash’ rejection (h - i). Figure objects: ‘lasso’ flagellum (red), posterior flagellum (blue), ingestion site (green circle), feeding current (solid curved arrows), object in motion (solid straight arrows).

### Particle tracks and clearance rates

For all species the particles followed hourglass-shaped paths, and only particles nearest the center of the flow were captured by the cell (Fig.6; Supplementary Fig. S2). To estimate clearance rates, two or three discs were placed for method validation but the estimated clearance rates remained almost the same within individuals (Supplementary Tab. S4). Estimated maximum clearance rates within species varied by a factor of about 2, and cell-volume specific clearance rates were all of the same order of magnitude, and varied from 2 – 18 × 10^6^ d^−1^ (Tab. 1).

**Figure 6.**
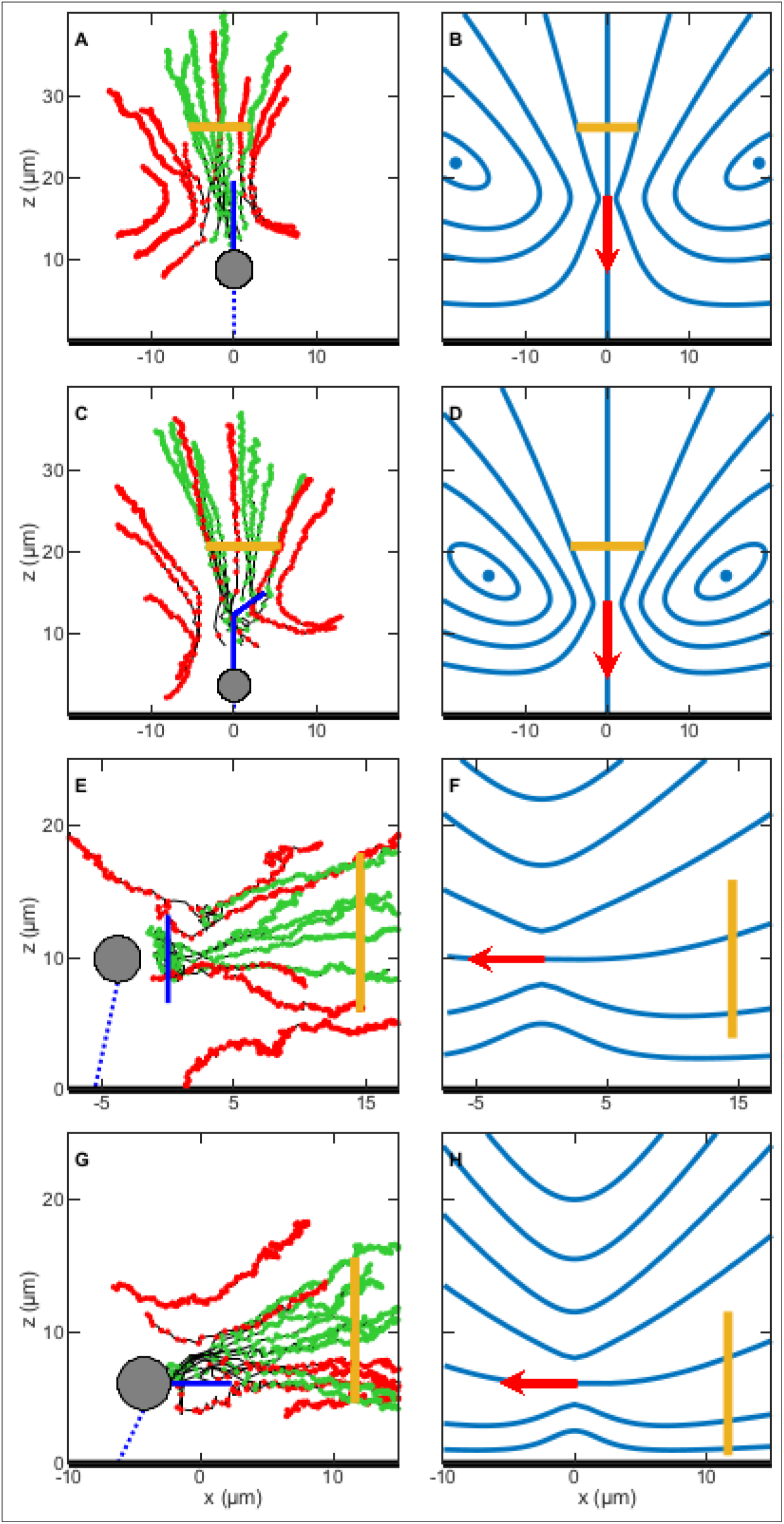
Observed feeding currents by particle tracking and theoretical flow fields generated with the point force model of the individuals Pteridomonas I (A and B), Paraphysomonas I (C and D), Pseudobodo I (E and F) and Cafeteria I (G and H). Clearance discs are represented as a yellow line. **Left-side panels:** the cell body (gray circle) of the flagellate attaches (blue dotted line) to the surface (thick black line), and the beating flagellum (solid blue line) generates a feeding current. Green tracks are for captured particles, and red tracks are for uncaptured prey. The positions of a particle, at each time-step (0.004 s), are represented as dots along the track. **Right-side panels:** the blue solid lines of the theoretical flows (B. D, F and H) depict the streamlines with equal flow rate, and the point force (red arrow) dictates the direction of the feeding current.

**Table 1.**
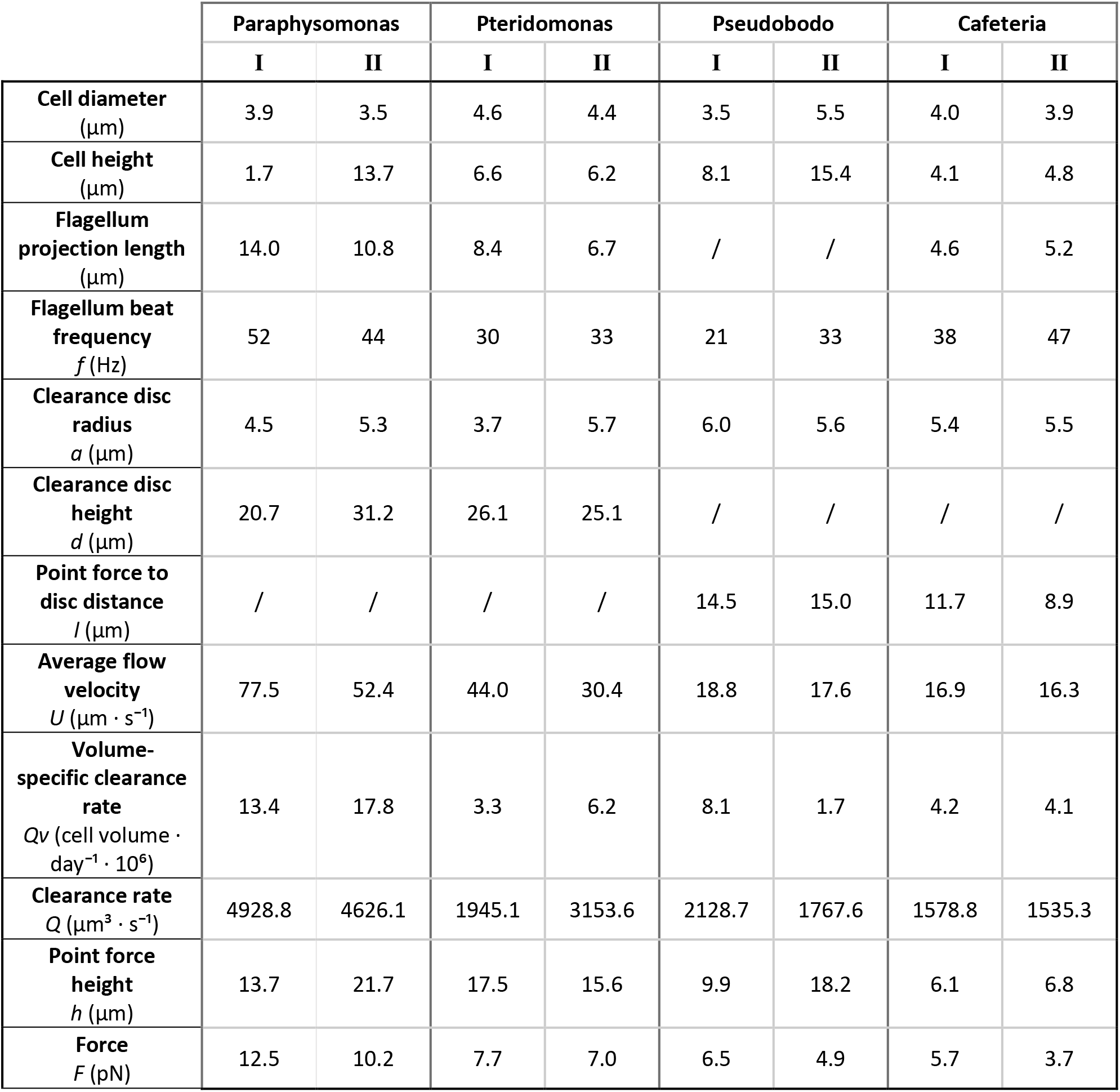
Clearance rates and force estimations of each study case. Particle tracking was performed to two individuals (I and II) of each species (*Paraphysomonas foraminifera*, *Pteridomonas danica*, *Pseudobodo* sp. and *Cafeteria roenbergensis*) to calculate the clearance rate and to estimate the force exerted by the flagellum. Relevant parameters to these results are included in the table.

### Estimation of force magnitudes and theoretical streamlines

The flow model assumes that the force acts at a *point*, while in reality the force production occurs *along* the flagellum. For *P. danica* we found the best fit for a point force located on the flagellum at 3/4 of its length (Fig.6A; Supplementary Fig. S2A). For *P. foraminifera* all particle tracks converged towards the curved distal segment of the flagellum of (Fig.6B; Supplementary Fig. S2B), therefore the point force was located in the middle of this section. Due to the complex and variable geometry of *Pseudobodo* sp., the point force was positioned in the middle of the ‘loop’ halfway between the body and the tip of the helical flagellum (Fig.6C; Supplementary Fig. S2C). The point force in the cases of *C. roenbergensis* was set to be at half the projection length of the flagellum, approximately where tracked particles reached maximum velocities (Fig.6D; Supplementary Fig. S2D). The resulting force estimates using equation (3) were 3.7 – 12.5 pN, with flows perpendicular to the surface (*P. danica* and *P. foraminifera*) requiring a slightly stronger force than the cases of parallel feeding currents (*Pseudobodo* sp. and *C. roenbergensis*) (Tab. 1).

The forces required to drive the observed flow can be compared with estimates of the force generated by the flagellum. *Pteridomonas danica* beats its flagellum in a plane with a roughly sinusoidal beat pattern, allowing us to apply the resistive force theory expression in equation (5) to directly estimate the force produced by the flagellum. We have our observed parameters *L* = 11.5 *μ*m, and 2 *A* = 2.9 *μ*m, *λ* = 5.3 *μ*m, and *f* = 33 Hz (Supplementary Tab. S1) and the parameters *b* = 0.15 *μ*m, *α* = 3 *μ*m, *β* = 0.01 *μ*m, and *χ* = 0.15 *μ*m from the literature [19, 33, 37]. With the parameters we would estimate *F_Z_* = −1.0 pN using equation (6) if the flagellum were without hairs, and we find *F_Z_* = 15 pN using the force coefficients in equation (7) for the flagellum with hairs. Similar estimates are not possible for the other species that have three-dimensional beat patterns.

## DISCUSSION

### Attached vs free-swimming

Our study complements earlier descriptions of the foraging behavior of heterotrophic nanoflagellates [e.g., 19, 25, 38]. We describe feeding only in flagellates attached to surfaces. Attached feeding appears to dominate for small forms, while larger forms, such as dinoflagellates, feed predominantly when free swimming [6, 39]. We observed *P. danica* capture prey while swimming, and *Spumella* sp., *O. moestrupii*, and loricated choanoflagellates are known to feed while swimming [18, 26, 30]. It has been argued that attachment enhances the feeding current of suspension feeders [13, 40, 41], but fluid dynamical simulations and models suggest the opposite [42, 43]. The reason for attachment in bacterivorous nanoflagellates may therefore rather be favorable food conditions near surfaces, such as marine snow [44, 45]. This is consistent with the observation that starving flagellates (*Ochromonas* sp, *P. vestita*, *A. mirabilis*) do not attach, while almost all cells experiencing high prey concentrations attach [13, 21]. Thus, swimming in heterotrophic nanoflagellates may for many species primarily serve the purpose of finding a nutrient-rich attachment surface. The probing behavior and different configuration of the flagellum of swimming and attached *Pseudobodo* sp. lend further support to this interpretation. Thus, stretching the flagellum in front of the cell allows faster swimming [46]. Some choanoflagellates similarly have an attached feeding stage with a short flagellum, and a free-swimming non-feeding stage with a long flagellum and a smaller more streamlined cell body [47]

### Handling time, clearance rate, and the functional response

Predator-prey interactions are often quantified by the prey ingestion rate as a function of the concentration of prey, typically described by a type II functional response [48]. This equation has two parameters, the maximum clearance rate, i.e., the volume of water cleared for particles per unit time at low prey concentration, and the prey handling time (= 1/maximum ingestion rate). Our behavioral observations allow us to estimate both parameters and to examine to what extent they underpin functional response relations estimated in incubation experiments.

In the species examined here, prey encounter is facilitated by the generation of a feeding current produced by the activity of one hairy flagellum that propels water towards the cell. We identified three different modes of prey encounter: prey particles arriving in the feeding current are either perceived and captured by the flagellum, intercepted by the cell body or by tentacles, and these represent the encounter mechanisms described for nanoflagellates. Prey is then handled by coordinated motions of one or two flagella, or, in the case of *P. danica*, by the tentacles. While the filter feeding *P. danica* continues to generate a feeding current while handling prey, the feeding current ceases during prey handling in the other species. The prey handling time can be substantial, particularly in *P. foraminifera* that stops beating the flagellum for up to more than 1 min after prey has been phagocytized. A similar’ refractory period’ has been reported for four species of nanoflagellates, including *C. roenbergensis* [25], leading to handling times between 4-95 s per prey, similar to the range reported here. Eventually, ingestion rate may be limited by handling times, and the so estimated maximum ingestion rates vary by more than one order of magnitude between species, from 1000 to 20,000 prey per day. This corresponds largely to the range of species-specific maximum ingestion rates of bacteria in incubation experiments, 600-6000 bacteria d^−1^ [6, 49]. The match becomes better when considering that a varying but sometimes large fraction of captured bacteria may be rejected [21, 24]. Handling of rejected prey may further reduce time for searching, even though handling time is generally shorter for rejected than ingested prey [6].

Particle tracking allowed us to characterize the flow field generated by the feeding flagellates, to identify the extension of the prey capture zone, and to estimate maximum clearance rates. Our estimates of cell-volume specific maximum clearance rates varied between both individuals and species, between 10^6^-10^7^ d^−1^. This magnitude is again similar to that obtained in incubation experiments, where estimates vary between species and range between 10^5^-10^7^ d^−1^ (reviewed in [8, 9]). Overall, functional responses measured in incubation experiments are mechanistically underpinned by behavioral observations.

### Flow architecture and fluid dynamics

At the low Reynolds number at which nanoflagellates operate, viscosity impedes predator-prey contact, but the activity of the beating flagellum is obviously sufficient to overcome the effect of viscosity. The impeding effect of viscosity is somewhat relaxed in flagellates that contact prey by the flagellum or tentacles at some distance from the no-slip surface of the cell. However, even in *C. roenbergensis*, where first contact is on the cell surface, the feeding current is sufficiently strong to allow prey encounters.

By applying a point force model that describes the observed flow fields well, we estimated the flow-generating forces to be on the order of 4-13 pN for the four species. These estimates ignore the presence of the cell body, and the force produced by the flagellum has to be somewhat larger than the force required to produce the observed flow fields. Christensen-Dalsgaard and Fenchel used an alternative approach and measured the swimming speed of *Paraphysomonas vestita* towing a latex sphere and computed the flagellum force from the Stokes drag to be of similar magnitude, 7-13 pN [19]. This approach neglects hydrodynamic interactions between flagellum, cell, and latex sphere, and the actual force is therefore larger than this estimate as well [46].

How do these indirect estimates compare with direct estimates of the force generated by the beating flagella? The estimate derived for *P. danica* by applying resistive force theory is larger but of similar magnitude as the indirect estimate, 15 pN and 7-8 pN, respectively. The estimate ignores hydrodynamic interactions between adjacent sections of the flagellum [42], which is most likely not justified for flagella with hairs, and it is therefore speculative. The estimate suggest that the hairs reverse the direction of the force and increases its magnitude by an order of magnitude compared to a flagellum without hairs. This increase is similar to that estimated by comparing swimming speeds of flagellates with smooth and hispid flagella [18].

As noted above, most heterotrophic nanoflagellates have hispid flagella, and this seems to be optimal or even necessary for prey encounter for a number of reasons. First, the presence of hairs significantly increases the force production of the beating flagellum and thereby the clearance rate. Secondly, the presence of hairs makes prey scanning of the flagellum efficient, since prey intercepted by the hairs elicits a capture response. Thirdly, the dominant flow along a flagellum with hairs is outside the envelope of the beating flagellum [50–52], presumably allowing efficient prey transport toward the cell. Forthly, the front-mounted flagellum increases the frequency of prey entrainment [53]. Finally, the reversal of the flow makes the streamlines come closer to the cell in the upstream direction from where the prey arrives, and the transport of captured prey towards the cell body is facilitated by the flow.

## Conclusions

Indirect and direct estimates of flagellum forces for one species are of similar magnitudes and consistent with the observed feeding flow, and the estimates of maximum ingestion and clearance rates are similar to those obtained from previous incubation experiments. Thus, our observations and estimates suggest a mechanistic underpinning of functional responses in heterotrophic nanoflagellates. However, experimentally estimated specific clearance rates of flagellates vary by two orders of magnitude [8, 9], and a significant fraction of this variation is accounted for by variation in flagellar arrangement and kinematics and consequent differences in flow architecture and predation risk: species with high clearance rates also disturb a large volume of water, attract flow-sensing predators from a further distance, and experience higher predation risk [18]. A better mechanistic understanding of this foraging trade-off and the variation in clearance rates requires a better understanding of the fluid dynamics of hairy flagella. This in turn may be facilitated by accurate observations of the often complex three-dimensional beat patterns of the flagella [14] and the arrangement of hairs on the flagella in combination with computational fluid dynamics simulations and theoretical modelling.

## MATERIALS AND METHODS

### Study organisms, isolation, and culturing

The four selected marine flagellate species *Pteridomonas danica, Paraphysomonas foraminifera, Cafeteria roenbergensis*, and *Pseudobodo* sp. are direct interception feeders (Fig. 1). They have a hairy (hispid) flagellum that drives the feeding current towards the cell, and in the opposite direction of the propagating flagellum wave. *Cafeteria roenbergensis* has been a key laboratory species, as its different feeding phases are easy to distinguish [20, 25]. Christensen-Dalsgaard and Fenchel explored the fluid dynamics of *P. danica* and *Paraphysomonas vestita*; the latter sharing the genus taxa with *P. foraminifera* [13, 19]. The feeding behavior of a close relative of *P. danica*, *Actinomonas* sp., has been studied [43, 25], and the predation mode of *Pseudobodo* sp. has been described briefly [19].A few observations of *Ochromonas moestrupii* and *Chrysophyceae* were also recorded.

All species were isolated from Danish waters. Cultures were maintained in the dark at 18°C in filtered and pasteurized North Sea water (salinity 30‰), using rice grains to feed the naturally occurring bacteria that served as prey. Species identification was verified using 18s rDNA molecular analysis except for *Pseudobodo* sp., which was morphologically matched with the description of *Pseudobodo tremulans* [24].

### Microscopy and image analysis

A glass ring (16 mm inner diameter and 3 mm height) was fixed on a glass slide using stopcock grease, filled with 600 μL of the culture, and covered with a glass coverslip. The sample was observed five minutes later to allow the flagellates to attach to the coverslip. The heating effects of light were reduced by having short periods of exposure (< 5 minutes) at moderate intensities during recordings. Room temperature was 16-20°C, and experiments did not last more than 1.5 hours. Food particles consisted of naturally occurring bacteria and particulate organic matter. For cultures with low bacterial abundance, polystyrene microbeads of 0.5 μm in diameter were added (10^−5^ %) to increase particle encounters.

Observations were carried out with an inverted microscope Olympus IX71, using an Olympus UPlanSApo oil immersion objective ×100 / 1.40 or an Olympus UPLanFL N oil immersion ×100 / 1.30 objective for phase contrast imaging. Recordings were carried out with a high-speed *Phantom Camera* (Miro LAB 320). Prey captures were recorded at 500 frames per second (fps), and particle tracking was performed at 250 fps. Videos had a minimum resolution of 512 × 512 pixels. The image analysis software ImageJ (Fiji) was used for detailed predator-prey interaction observations, morphometric measurements, and manual particle tracking [31, 32].

### Predation behavior and time budget analysis

The foraging behavior was observed in different individuals of the same exponentially- grown young culture of each species (biological replicates). Predation was divided in five stages, largely following [26]: searching, contact, capture, ingestion or rejection, and recovery. A rejection was defined as an active release of the prey, in contrast to a loss of prey. The handling time was defined as the duration of the period where the flagellum produces no feeding current and another prey cannot be encountered. Handling time does not necessarily commence upon prey contact, and it can end before or after the particle is fully ingested. Prey processing had three possible outcomes: ingestion, rejection, or loss. In total, 40 prey handling durations (20 ingestions and 20 rejections) were recorded for each species. In addition, 20 ‘lash rejections’ by *Pseudobodo* sp. were analyzed (Supplementary Tab. S2). This research is based on behavior observations and is a non-interventional study. Thus, considering the nature of the analysis, the sample sizes N = 40 – 60 were regarded as satisfactory.

### Particle tracking and clearance rates

Flow fields were mapped by particle tracking. A particle was followed from a minimum distance of one cell length from the body, until it was either captured or it had gone well past the flagellate. We recorded 11 – 15 tracks per individual, studying two individuals per species. Most flagellates slightly shift their orientation while foraging (Supplementary Tab. S3), and the particle tracks are therefore shown relative to the observed flagellum coordinates. An imaginary, circular filtering area (clearance disc) for prey capture was assumed in front of the cell and perpendicular to the feeding flow. The size of the disc was defined by the tracks of particles that were captured or strongly interacted with the flagellate. A five-point, centered finite difference scheme was applied to the measured particle positions to minimize noise and discretization errors when calculating the particle velocities. The average velocity component perpendicular to the clearance disc was used to calculate the clearance rate.

### Model of the feeding flow

To describe the flow fields and estimate the flow-generating forces from the observed feeding currents we use a point force model [12, 27–29]. The low-Reynolds-number model describes the flow due to a point force above a plane no-slip surface. We examined two situations in which the force direction is either perpendicular (*P. danica*, *P. foraminifera*) or parallel to the surface (*Pseudobodo* sp., *C. roenbergensis*). In both cases, we use *F* to denote the magnitude of the force and *h* for its height above the surface. In the perpendicular case, the flow has rotational symmetry and the streamlines are the contour lines of the Stokes stream function:

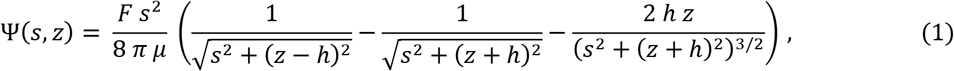

where *μ* denotes the viscosity, *s* the radial distance from the axis of symmetry, and *z* the height above the surface [30, 31]. Using the stream function, we can derive the clearance rate,*Q*, through a circular clearance disc centered on the axis of symmetry and oriented perpendicular to it:

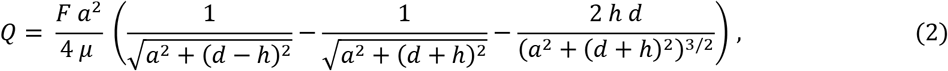

where *a* denotes the radius of the clearance disc and *d* its height above the surface [29]. The equation allows us to estimate the magnitude of the point force,*F*, using our clearance rate estimate obtained with particle tracking. In the parallel case, the flow does not have rotational symmetry and a Stokes stream function does not exist. We therefore integrate the velocity field numerically to obtain streamlines [28, 29]. To estimate the clearance rate through a circular clearance disc that is perpendicular to the direction of the point force and positioned the distance *h* above the surface in the symmetry plane of the flow, we assume that *a* ≪ *h* and approximate the effect of the image system that ensures that the no-slip boundary condition is satisfied [29]. We find the approximation:

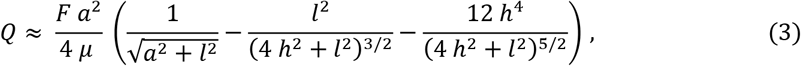

where *l* denotes the distance between the position of the point force and the center of the clearance disc.

### The force created by a hairy flagellum

To estimate the flow-generating force of *P. danica* directly from the observed motion of the flagellum, we use resistive force theory [32–36]. We assume that the flagellate is tethered, and that the motion of the flagellum is a travelling sine wave with wavelength λ and frequency *f* that is propagating in the positive *z*-direction:

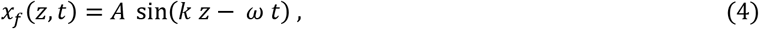

where *x_f_* denotes the transversal deflection of the flagellum, *A* the amplitude, *k* = 2 *π/λ* the wave number, and *ω* = 2 π *f* the angular frequency. Using resistive force theory, we obtain the component of the force in the *z*-direction:

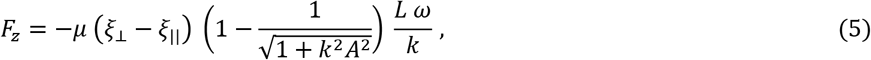

where *L* denotes the length of the flagellum, and ξ_⊥_and ξ_||_ the perpendicular and the parallel force coefficient, respectively [41]. For a naked flagellum we use the dimensionless force coefficients:

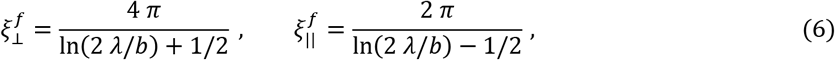

where *b* denotes the radius of the flagellum [32, 36]. For a flagellum with two rows of stiff hairs that are in the beat plane and remain perpendicular to the flagellum during the beat, we use the dimensionless force coefficients:

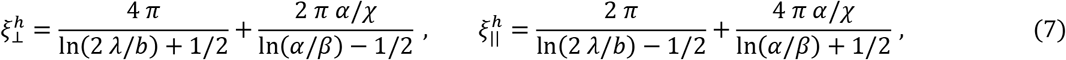

where *α* denotes the total length of a pair of hairs, *β* their radius, and *χ* the distance between neighboring pairs of hairs [33, 34].

## Acknowledgements

We acknowledge help from Ramon Massana, Vanessa Balagué, Elisabet Laia Sà, Lasse Nielsen, Helge Thomsen, and Mads Rode. We received funding from The Independent Research Fund Denmark (7014-00033B) and the Carlsberg Foundation (CF17-0495). The Centre for Ocean Life is supported by the Villum Foundation,).

## SUPPLEMENTARY INFORMATION

**Supplementary Figure S1.**
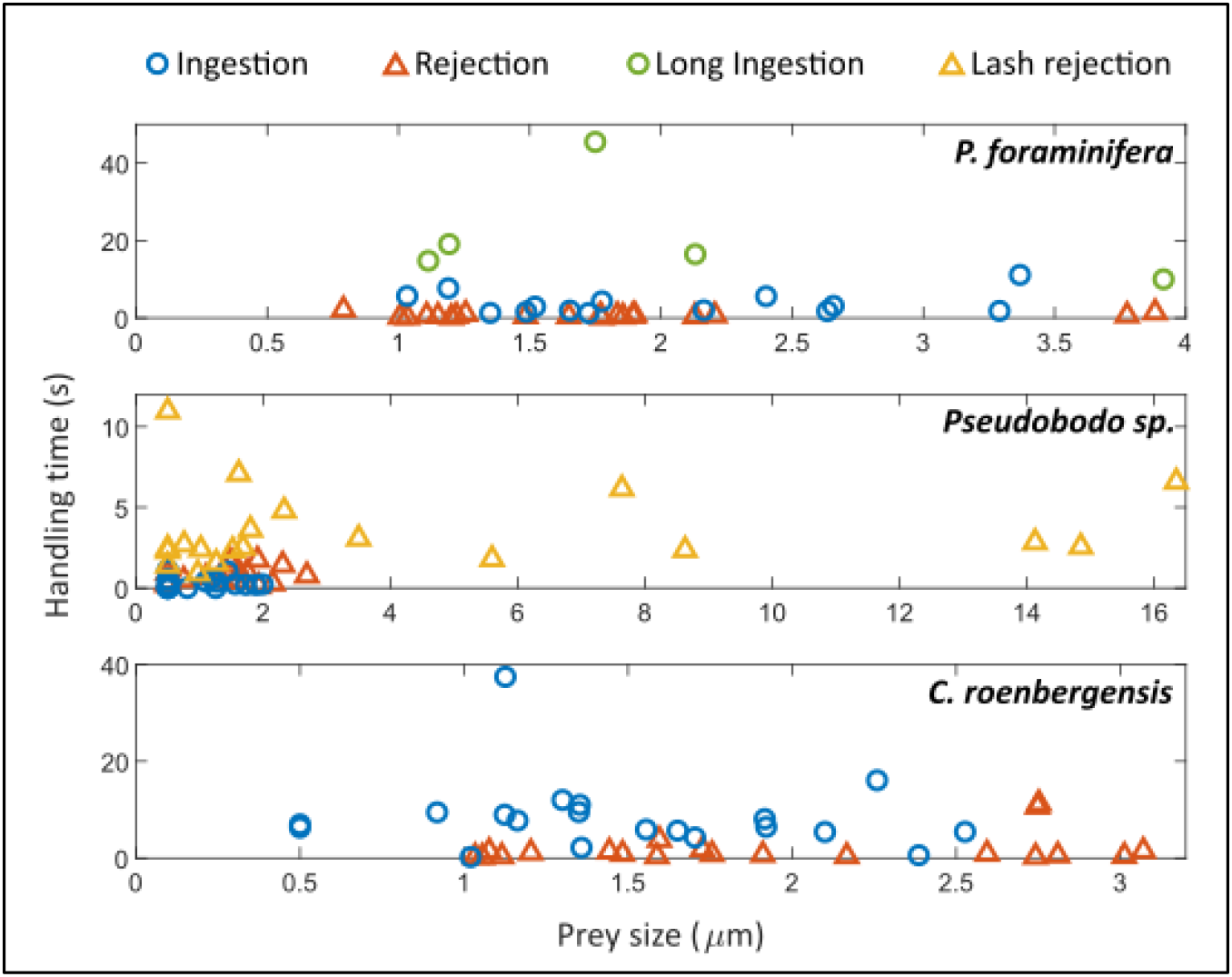
Handling times for different prey sizes (when available) by *Paraphysomonas foraminifera*, *Pseudobodo* sp., and *Cafeteria roenbergensis*. No correlation was found between the duration of prey handling (for ingestions and rejections) and the prey size in all studied species. For *P. foraminifera*, ‘Long ingestions’ (N = 4) are the minimum handling times that were limited by the recording capacity (i.e. when the flagellum stopped beating).

**Supplementary Figure S2.**
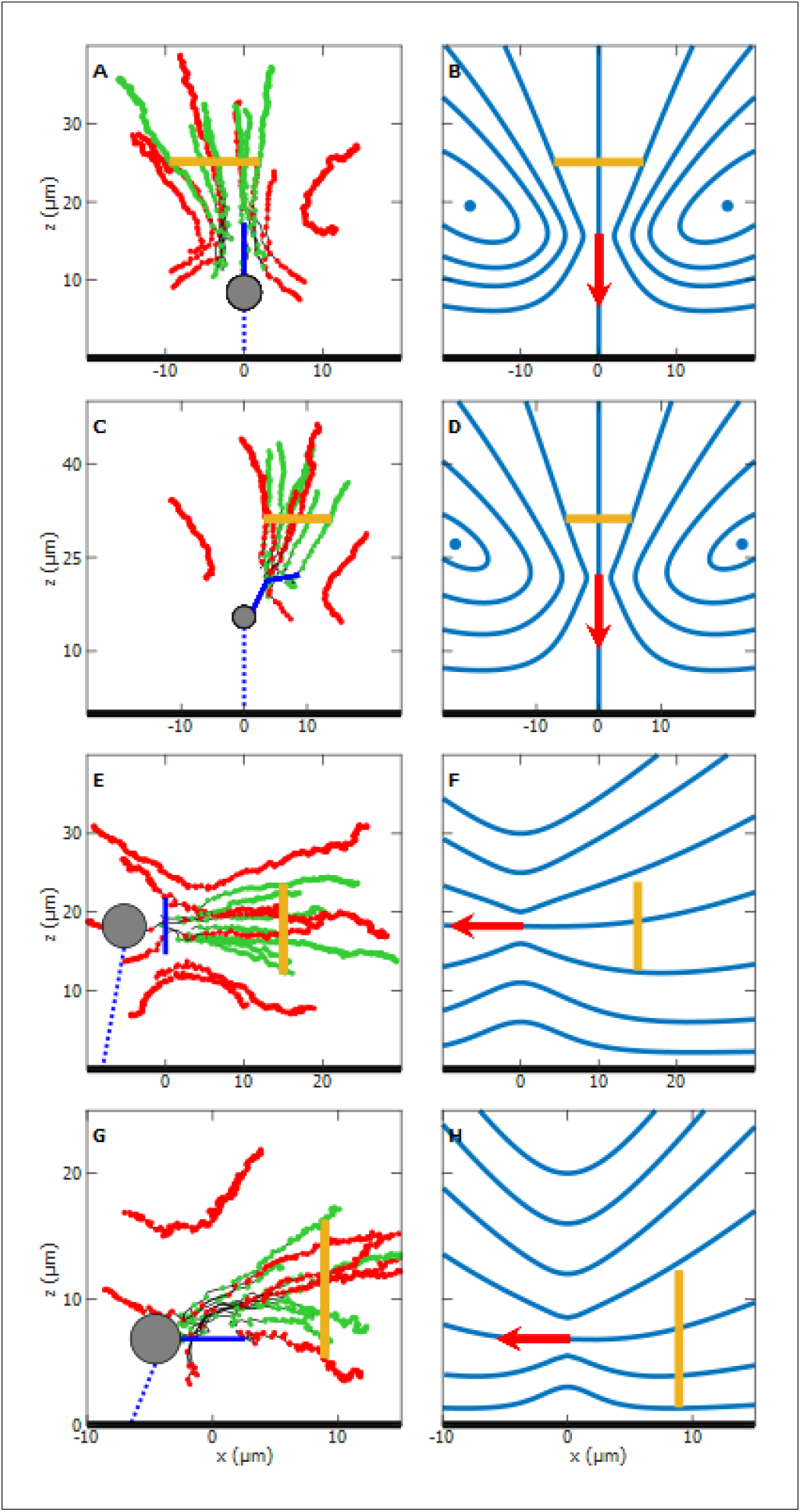
Observed feeding currents by particle tracking and theoretical flow fields generated with the point force model of the individuals Pteridomonas II (A and B), Paraphysomonas II (C and D), Pseudobodo II (E and F) and Cafeteria II (G and H). Clearance discs are represented as a yellow line. **Left-side panels:** the cell body (gray circle) of the flagellate attaches (blue dotted line) to the surface (thick black line), and the beating flagellum (solid blue line) generates a feeding current. Green tracks are for captured particles, and red tracks are for uncaptured prey. The positions of a particle, at each time-step (0.004 s), are represented as dots along the track. **Right-side panels:** the blue solid lines of the theoretical flows (B. D, F and H) depict the streamlines with equal flow rate, and the point force (red arrow) dictates the direction of the feeding current.

**Supplementary Table S1.**
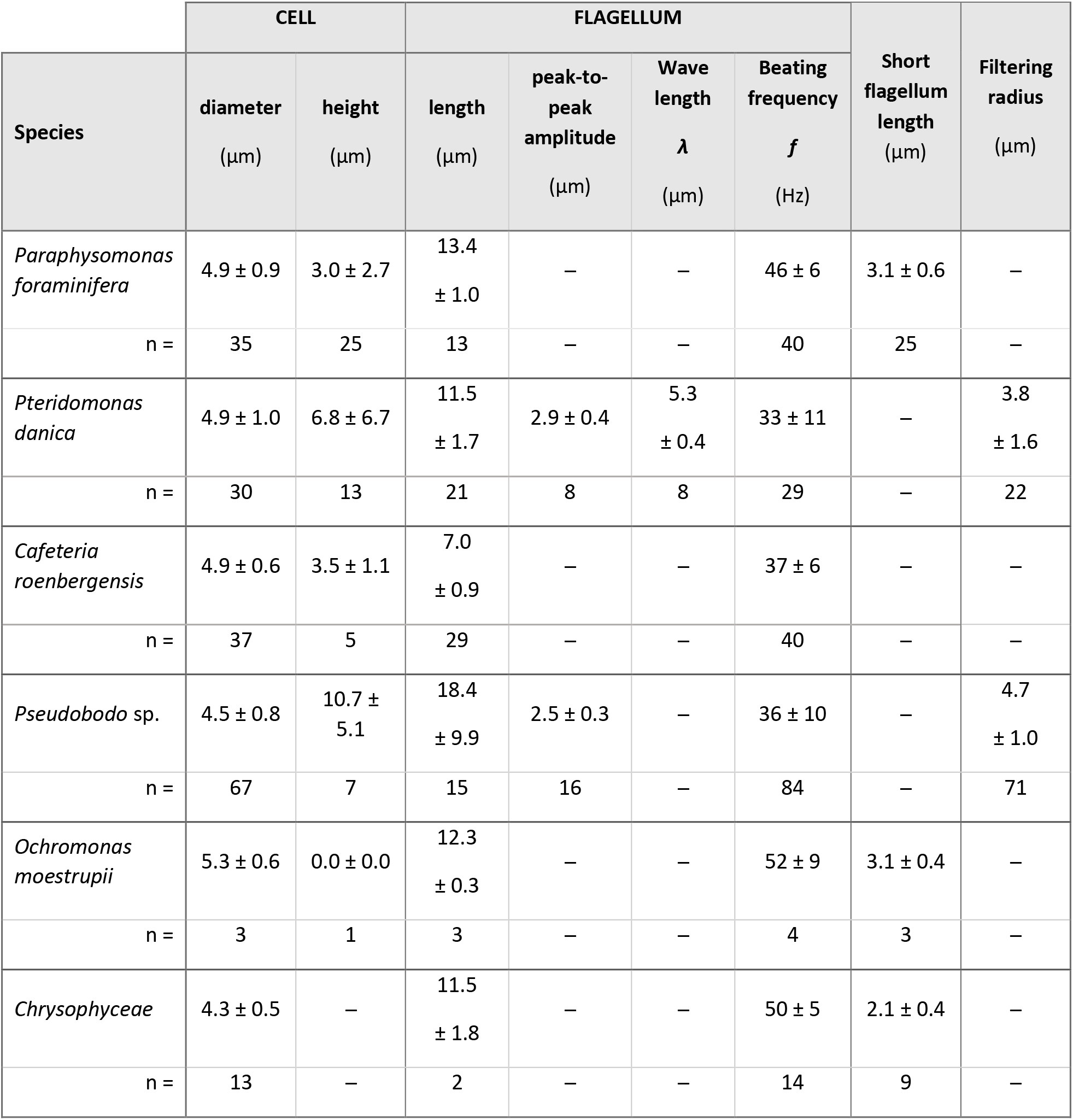
Morphometric data and flagellum properties. Cell height is the distance between the substrate to the attached cell. Flagellum features correspond to flagella creating a feeding current, i.e. not interacting with prey. The flagellum amplitude was measured from peak-to-peak of the wave (note that for *Pseudobodo* sp. it is the length of the 2D projection from the 3D beating wave). Short flagellum refers to the second prey-handling flagellum of *P. foraminifera*, *O. moestrupii* and *Chrysophyceae*. The filtering radius is the distance from the tip of a lateral tentacle to the central axis of the body or the radius of the flagellar ‘loop’ for *P. danica* and *Pseudobodo* sp., respectively. The measurements are expressed as the mean ± standard deviation, and the number of observations per parameter is N.

**Supplementary Table S2.** Handling times during ingestions and rejections. * Not including the cases (N = 5) when *P. foraminifera* stops beating during ingestions that were limited by the recording time.

| Species | Handling Time (s) |  |  |  |  |  |
| --- | --- | --- | --- | --- | --- | --- |
|  | Ingestion |  | Rejection |  | Lash Rejection |  |
| | N | mean ± SD | N | mean ± SD | N | mean ±SD |
| *Pteridomonas danica* | 0 |  | 0 |  | – |  |
| *Paraphysomonas foraminifera* | 15* | 5.7 ± 7.8* | 20 | 0.6 ± 0.5 | – | |
| *Pseudobodo* sp. | 20 | 0.3 ± 0.3 | 20 | 0.7 ± 0.5 | 20 | 3.5 ± 2.4 |
| *Cafeteria roenbergensis* | 20 | 8.5 ± 7.6 | 20 | 1.9 ± 3.1 | – | |
| *Ochromonas moestrupii* | 1 | 5.5 | 2 | 1.2 ± 0.8 | – | |
| *Chrysophyceae* | – | | 12 | 5.2 ± 6.0 | – | |

**Supplementary Table S3.** Tilting angle (α) of the force vector generated by the beating flagellum. Minor shifts of orientation while generating feeding currents were observed in *Pteridomonas danica*, *Paraphysomonas foraminifera*, and *Pseudobodo* sp. Angle α lays between the force vector and the axis normal to the surface. The force is orthogonal to the surface when α = 0°, and parallel when α = 90°. Alpha was recorded in each track for later averaging.

| Species individual | Number of Tracks | Average $\alpha$ (°) | SD |
| --- | --- | --- | --- |
| Pteridomonas I | 15 | 7.3 | 3.9 |
| Pteridomonas II | 15 | 10.7 | 3 |
| Paraphysomonas I | 15 | 14.1 | 5.3 |
| Paraphysomonas II | 13 | 8.7 | 3.9 |
| Pseudobodo I | 11 | 57.2 | 9.3 |
| Pseudobodo II | 14 | 92.5 | 10.2 |

**Supplementary Table S4.** Clearance rates (Q) from each disc of radius a, placed at different distances (distance cell to disc) from the species individual. As a method validation, the flow rate through more than one clearance disc per individual was estimated and compared. Variations within each species (*Paraphysomonas foraminifera*, *Pteridomonas danica*, *Pseudobodo* sp. and *Cafeteria roenbergensis*) are by a factor of around two.

| | | Clearance disc radius<br>$a$ ($\mu\text{m}$) | Distance cell to disc<br>($\mu\text{m}$) | Clearance rate<br>$Q$ ($\mu\text{m}^3 \cdot \text{s}^{-1}$) |
| --- | --- | --- | --- | --- |
| **Pteridomonas I** | Disc 1 | 2.3 | 10.0 | 1374.4 |
|  | Disc 2 | 3.8 | 15.0 | 1945.1 |
|  | Disc 3 | 5.5 | 20.0 | 2212.6 |
| **Pteridomonas II** | Disc 1 | 3.8 | 10 | 2672.4 |
|  | Disc 2 | 5.8 | 14.5 | 3153.6 |
|  | Disc 3 | 7.5 | 19 | 3464.9 |
| **Paraphysomonas I** | Disc 1 | 3.0 | 10.0 | 4503.8 |
|  | Disc 2 | 4.5 | 15.0 | 4928.8 |
|  | Disc 3 | 6.3 | 20.0 | 5511.2 |
| **Paraphysomonas II** | Disc 1 | 4.2 | 10.0 | 5029.2 |
|  | Disc 2 | 5.3 | 14.0 | 4626.1 |
|  | Disc 3 | 6.3 | 18.0 | 4565.8 |
| **Pseudobodo I** | Disc 1 | 5.0 | 11.5 | 1836.8 |
|  | Disc 2 | 6.0 | 16.5 | 2128.7 |
| **Pseudobodo II** | Disc 1 | 4.2 | 12.5 | 1909.5 |
|  | Disc 2 | 5.7 | 17.5 | 1767.6 |
| **Cafeteria I** | Disc 1 | 2.4 | 5.0 | 1454.6 |
|  | Disc 2 | 3.9 | 10.0 | 1128.4 |
|  | Disc 3 | 5.5 | 14.0 | 1578.8 |
| **Cafeteria II** | Disc 1 | 2.5 | 5.5 | 1473.5 |
|  | Disc 2 | 4.1 | 8.5 | 1888.0 |
|  | Disc 3 | 5.5 | 11.5 | 1535.3 |

## REFERENCES

1. Azam F, Fenchel T, Field J, Gray J, Meyer-Reil L, Thingstad TF. The Ecological Role of Water-Column Microbes in the Sea. Mar Ecol Prog Ser 1983; 10: 257–263.

2. Worden AZ, Follows MJ, Giovannoni SJ, Wilken S, Zimmerman AE, Keeling PJ. Rethinking the marine carbon cycle: Factoring in the multifarious lifestyles of microbes. Science (80-) 2015; 347: 1257594–1257594.

3. Fenchel T. Ecology of Heterotrophic Microflagellates. IV Quantitative Occurrence and Importance as Bacterial Consumers. Mar Ecol Prog Ser 1982; 9: 35–42.

4. Jürgens K, Matz C. Predation as a shaping force for the phenotypic and genotypic composition of planktonic bacteria. Antonie van Leeuwenhoek, Int J Gen Mol Microbiol 2002; 81: 413–434.

5. Weisse T, Anderson R, Arndt H, Calbet A, Hansen PJ, Montagnes DJS. Functional ecology of aquatic phagotrophic protists – Concepts, limitations, and perspectives. Eur J Protistol 2016; 55: 50–74.

6. Boenigk J, Arndt H. Bacterivory by heterotrophic flagellates: community structure and feeding strategies. Antonie Van Leeuwenhoek 2002; 81: 465–480.

7. Jabbarzadeh M, Fu HC. Viscous constraints on microorganism approach and interaction. J Fluid Mech 2018; 851: 715–738.

8. Hansen PJ, Bjørnsen PK, Hansen BW. Zooplankton grazing and growth: Scaling within the 2-2,-μm body size range. Limnol Oceanogr 1997; 42: 687–704.

9. Kiørboe T, Hirst AG. Shifts in mass scaling of respiration, feeding, and growth rates across life-form transitions in marine pelagic organisms. Am Nat 2014; 183: E118–30.

10. Kiørboe T. How zooplankton feed: Mechanisms, traits and trade-offs. Biol Rev 2011; 86: 311–339.

11. Fenchel T. Suspended marine bacteria as a food source. Suspended marine bacteria as a food source, In: Fasham, M. J. (ed.), Flow of Material and Energy in Marine Ecosystems. Plenum Press, New York. 1984. pp 301–315.

12. Fenchel T. Protozoan filter feeding. Prog Prostit 1986; 1: 65–113.

13. Christensen-Dalsgaard K, Fenchel T. Increased filtration efficiency of attached compared to free-swimming flagellates. Aquat Microb Ecol 2003; 33: 77–86.

14. Christensen-Dalsgaard KK, Fenchel T. Complex Flagellar Motions and Swimming Patterns of the Flagellates and. Protist 2004; 155: 79–87.

15. Dölger J, Nielsen LT, Kiørboe T, Andersen A. Swimming and feeding of mixotrophic biflagellates. Sci Rep 2017; 7: 39892.

16. Nielsen LT, Asadzadeh SS, Dölger J, Walther JH, Kiørboe T, Andersen A. Hydrodynamics of microbial filter feeding. Proc Natl Acad Sci U S A 2017; 114.

17. Roper M, Dayel MJ, Pepper RE, Koehl MAR. Cooperatively Generated Stresslet Flows Supply Fresh Fluid to Multicellular Choanoflagellate Colonies. Phys Rev Lett 2013; 110: 228104.

18. Nielsen LT, Kiørboe T. Foraging trade-offs, flagellar arrangements, and flow architecture of planktonic protists. Proc Natl Acad Sci 2021; 118: e2009930118.

19. Fenchel T. Ecology of Heterotrophic Microflagellates. I. Some Important Forms and Their Functional Morphology. Mar Ecol Prog Ser 1982; 8: 211–223.

20. Ishigaki T, Terazaki M. Grazing behavior of heterotrophic nanoflagellates observed with a high speed VTR system. J Eukaryot Microbiol 1998; 45: 484–487.

21. Pfandl K, Posch T, Boenigk J. Unexpected effects of prey dimensions and morphologies on the size selective feeding by two bacterivorous flagellates (Ochromonas sp. and Spumella sp.). J Eukaryot Microbiol 2004; 51: 626–633.

22. Boenigk J, Arndt H. Comparative studies on the feeding behavior of two heterotrophic nanoflagellates: The filter-feeding choanoflagellate Monosiga ovata and the raptorial-feeding kinetoplastid Rhynchomonas nasuta. Aquat Microb Ecol 2000; 22: 243–249.

23. Wu QL, Boenigk J, Hahn MW. Successful Predation of Filamentous Bacteria by a Nanoflagellate Challenges Current Models of Flagellate Bacterivory. Appl Environ Microbiol 2004; 70: 332–339.

24. Matz C, Boenigk J, Arndt H, Jürgens K. Role of bacterial phenotypic traits in selective feeding of the heterotrophic nanoflagellate Spumella sp. Aquat Microb Ecol 2002; 27: 137–148.

25. Boenigk J, Arndt H. Particle handling during interception feeding by four species of heterotrophic nanoflagellates. J Eukaryot Microbiol 2000; 47: 350–358.

26. Montagnes DJS, Barbosa AB, Boenigk J, Davidson K, Jürgens K, Macek M, et al. Selective feeding behaviour of key free-living protists: Avenues for continued study. Aquat Microb Ecol 2008; 53: 83–98.

27. Blake JR. A note on the image system for a stokeslet in a no-slip boundary. Math Proc Cambridge Philos Soc 1971; 70: 303–310.

28. Blake JR, Chwang AT. Fundamental singularities of viscous flow. J Eng Math 1974; 8: 23–29.

29. Rode M, Meucci G, Seegert K, Kiørboe T, Andersen A. Effects of surface proximity and force orientation on the feeding flows of microorganisms on solid surfaces. Phys Rev Fluids 2020; 5: 123104.

30. Aderogba K, Blake JR. Action of a force near the planar surface between semi-infinite immiscible liquids at very low Reynolds numbers: Addendum. Bull Aust Math Soc 1978; 19: 309–318.

31. Blake JR, Otto SR. Ciliary propulsion, chaotic filtration and a blinking stokeslet. J Eng Math 1996; 151–168.

32. Gray J, Hancock GJ. The propulsion of sea-urchin spermatozoa. J Exp Biol 1955; 32: 802–814.

33. Holwill MA, Sleigh M. Propulsion by hispid flagella. J Exp Biol 1967; 47: 267–276.

34. Brennen C. Locomotion of flagellates with mastigonemes. J Mechanochemistry Cell Motil 1976; 3: 207–217.

35. Lauga E, Powers TR. The hydrodynamics of swimming microorganisms. Reports Prog Phys 2009; 72: 096601.

36. Rodenborn B, Chen C-H, Swinney HL, Liu B, Zhang HP. Propulsion of microorganisms by a helical flagellum. Proc Natl Acad Sci 2013; 110: E338–E347.

37. Patterson DJ, Fenchel T. Insights into the evolution of heliozoa (Protozoa, Sarcodina) as provided by ultrastructural studies on a new species of flagellate from the genus Pteridomonas. Biol J Linn Soc 1985; 34: 381–403.

38. Sleigh BMA. Flagellar movement of the Sessile Flagellates Actinomonas, Codonosiga, Monas, and Poteriodendron. J Cell Sci 1964; s3-105: 405–414.

39. Nielsen LT, Kiørboe T. Feeding currents facilitate a mixotrophic way of life. ISME J 2015; 9: 2117–2127.

40. Strickler JR. Calanoid Copepods, Feeding Currents, and the Role of Gravity. Science (80-) 1982; 218: 158–160.

41. Tiselius P, Jonsson P. Foraging behaviour of six calanoid copepods: observations and hydrodynamic analysis. Mar Ecol Prog Ser 1990; 66: 23–33.

42. Kirkegaard JB, Goldstein RE. Filter-feeding, near-field flows, and the morphologies of colonial choanoflagellates. Phys Rev E 2016; 94: 052401.

43. Andersen A, Kiørboe T. The effect of tethering on the clearance rate of suspension-feeding plankton. Proc Natl Acad Sci 2020; 202017441.

44. Alldredge AL, Silver MW. Characteristics, dynamics and significance of marine snow. Prog Oceanogr 1988; 20: 41–82.

45. Simon M, Grossart HP, Schweitzer B, Ploug H. Microbial ecology of organic aggregates in aquatic ecosystems. Aquat Microb Ecol 2002; 28: 175–211.

46. Langlois V, Andersen A, Bohr T, Visser A, Kiørboe T. Significance of swimming and feeding currents for nutrient uptake in osmotrophic and interception feeding flagellates. Aquat Microb Ecol 2009; 54: 35–44.

47. Nguyen H, Koehl MAR, Oakes C, Bustamante G, Fauci L. Effects of cell morphology and attachment to a surface on the hydrodynamic performance of unicellular choanoflagellates. J R Soc Interface 2019; 16: 20180736.

48. Holling CS. Some Characteristics of Simple Types of Predation and Parasitism. Can Entomol 1959; 91: 385– 398.

49. Fenchel T. Ecology of Heterotrophic Microflagellates. II. Bioenergetics and Growth. Mar Ecol Prog Ser 1982; 8: 225–231.

50. Jahn TL, Landman MD, Fonseca JR. The Mechanism of Locomotion of Flagellates. II. Function of the Mastigonemes of Ochromonas. J Protozool 1964; 3: 291–296.

51. Sleigh M a. Flagellar beat patterns and their possible evolution. Biosystems 1981; 14: 423–431.

52. Sleigh MA. Mechanisms of flagellar propulsion - A biologist’s view of the relation between structure, motion, and fluid mechanics. Protoplasma 1991; 164: 45–53.

53. Mathijssen AJTM, Jeanneret R, Polin M. Universal entrainment mechanism controls contact times with motile cells. Phys Rev Fluids 2018; 3: 033103.

